# High-fat diet promotes exhaustion-associated T-cell remodeling across vascular and lymphoid tissues in atherosclerosis-prone mice

**DOI:** 10.64898/2026.11.03.763491

**Authors:** Pourya Yarahmadi, Yinghua Xie, Diem Thi Ngoc Huynh, Jacquelyn Lai, Qianhui Ling, Xianxi Huang, Patricia K. Nguyen

## Abstract

**Background:** Atherosclerosis is a chronic inflammatory disease driven by lipid accumulation and immune dysregulation. How chronic dietary lipid exposure reshapes T-cell states across vascular and lymphoid tissues, and whether these changes resolve after dietary normalization, remains incompletely defined. This study examined diet-associated T-cell remodeling and the contribution of macrophage-mediated stimulation

**Methods:** ApoE-deficient mice received a vegetarian diet (VD) or high-fat diet (HFD) for 12 or 24 weeks; a dietary reversal group received HFD for 12 weeks followed by VD for 12 weeks. CD45-positive cells from aorta, blood, spleen, lymph nodes, and bone marrow underwent single-cell RNA sequencing with matched T-cell receptor sequencing. T-cell states were annotated using canonical markers, and exhaustion-and naive-associated transcriptional programs were quantified using UCell scores. Analyses assessed T-cell composition, clonal expansion, and predicted macrophage–T-cell communication, with splenic flow-cytometric validation.

**Results:** HFD was associated with enrichment of exhaustion-associated states, with the highest exhaustion scores in the aorta. At 24 weeks, the pooled exhausted-cell proportion was 15.7% under HFD versus 10.6% under VD. Dietary reversal restored this proportion to 10.6%, while residual transcriptional changes persisted in a tissue-dependent manner. Flow cytometry corroborated increased inhibitory receptor expression and co-expression. Aortic macrophages exhibited inflammatory and antigen-presentation programs alongside increased predicted communication with T cells. Aortic T-cell activation and clonal expansion accompanied these changes, and clonotype size correlated with exhaustion score (Spearman correlation coefficient = 0.57, p = 0.029).

**Conclusion:** Chronic HFD is associated with multicompartment T-cell remodeling, with prominent vascular exhaustion-associated programs and tissue-dependent recovery after dietary normalization. These findings suggest that dietary lipid exposure may leave a durable adaptive immune imprint across organs during atherosclerosis.

## Introduction

Atherosclerosis is now understood as a chronic inflammatory disease driven by the retention of lipoproteins within the arterial wall and by coordinated innate and adaptive immune responses that shape lesion progression and instability.^1,2^ Although macrophages are central to plaque development, T cells are abundant in atherosclerotic lesions and increasingly recognized as important regulators of vascular inflammation, plaque composition, and clinical outcomes.^3,4^ These observations support the view that atherosclerosis is not merely a disorder of lipid deposition, but a systemic immune-metabolic disease linking circulating lipids, lymphoid organs, and the vessel wall.

Hyperlipidemia has the potential to influence this network at multiple levels. Cholesterol is not only a metabolic substrate but also an essential structural component of cellular membranes that can modulate T-cell receptor organization, signaling, and cell fate. In experimental systems, hypercholesterolemia enhances T-cell receptor signaling, and cholesterol-rich environments can drive inhibitory receptor expression and exhaustion-like dysfunction in T cells.^5,6^ These findings raise the possibility that chronic dietary lipid exposure may progressively reprogram T-cell state both systemically and within vascular tissues.

This question is especially relevant in atherosclerosis, where persistent antigen exposure and repeated antigen-presenting-cell–T-cell interactions may favor chronic stimulation. ApoB-reactive CD4+ T cells have emerged as key participants in disease, with studies in mice and humans demonstrating clonal expansion, phenotypic plasticity, and progressive conversion toward pathogenic states during atherogenesis.^3,7^ In parallel, single-cell analyses of human atherosclerotic plaques have revealed marked immune heterogeneity, including activated and checkpoint-rich T-cell states.^8-10^ More recent work further suggests that progenitor-exhausted-like PD-1+ T cells may represent an important vascular T-cell population in atherosclerosis, linking exhaustion-associated biology directly to vascular disease.^11^

Despite these advances, most single-cell studies of atherosclerosis remain cross-sectional and plaque-centric, leaving unresolved how chronic dietary lipid exposure reshapes T-cell state over time across vascular and lymphoid compartments. It is also unclear whether these changes are also reversible when lipid exposure is reduced and how local macrophage–T-cell interactions contribute to vascular T-cell dysfunction. Here, we used longitudinal multi-tissue single-cell RNA sequencing to define how chronic high-fat diet remodels T-cell states across the aorta, spleen, lymph nodes, bone marrow, and blood. We show that chronic dietary lipid exposure progressively shifts T cells toward exhaustion-associated programs across tissues, with the strongest burden in the aorta, and that this aortic phenotype is associated with inflammatory, antigen-presenting macrophages, enhanced macrophage-to-T-cell communication, heightened T-cell activation, and clonal expansion.

## Methods

### Animals, diets, and study design

All animal procedures were performed in accordance with institutional guidelines and approved protocols. Unless otherwise indicated, experiments were conducted using female mice. For atherosclerosis and multi-tissue immune profiling, ApoE□/□ mice were maintained on either a vegetarian control diet (VD; Teklad 2918; energy density 3.1 kcal/g; manufacturer-reported macronutrient composition with no added cholesterol) or a cholesterol- and fat-enriched high-fat diet (HFD; Teklad TD.88137; energy density 4.5 kcal/g; ∼42% kcal from fat; manufacturer-reported cholesterol 2076.6 mg/kg). Diets were initiated at 6 weeks of age (baseline), and mice were analyzed at baseline or after 12 or 24 weeks of feeding. For dietary normalization (reversed diet, RD), mice were maintained on HFD for 12 weeks and then switched to VD for another 12 weeks prior to analysis. At designated time points, mice were euthanized and tissues were harvested for downstream analyses, including aorta, spleen, lymph nodes, bone marrow, and peripheral blood (PBMCs).

### Tissue processing and single-cell library generation

At the experimental endpoint, mice were euthanized and tissues were harvested into cold buffer for immediate processing. To reduce contamination by circulating blood, mice were perfused before vascular tissue isolation. Aortas were dissected, cleaned, minced, and enzymatically digested to generate single-cell suspensions. Spleens and lymph nodes were mechanically dissociated, bone marrow was flushed from long bones, and peripheral blood mononuclear cells were isolated by density-gradient centrifugation. For scRNA-seq, viable CD45+ immune cells were isolated by fluorescence-activated cell sorting. Single-cell gene-expression libraries were generated using the 10x Genomics Chromium 5′ platform, and matched TCR V(D)J libraries were prepared from the same partitions.

### Single-cell RNA-seq analysis

Raw sequencing data were processed to generate filtered gene-barcode matrices and annotated TCR contigs. After standard quality control and removal of low-quality cells and likely doublets, datasets were normalized, integrated across samples, and clustered. Major immune populations were first identified in the global CD45+ dataset, after which T cells were subsetted and reanalyzed to improve lymphoid resolution. Final T-cell annotations were assigned using iterative clustering together with manual curation based on canonical markers and cluster-level differential expression. The broad T-cell states used in the main figures were naive/central memory T cells, effector memory/effector T cells, exhausted T cells, regulatory T cells, γδ T cells, other T helper cells, and progenitor/proliferating T cells. Differential expression between diet groups and time points was then used to identify transcriptional programs associated with chronic HFD exposure and dietary reversal.

### Quantification of T-cell composition

T-cell composition was summarized in two ways. For descriptive stacked-bar plots, annotated T-cell states were pooled within each group or tissue-by-group subset and displayed as relative proportions. For statistical comparisons, proportions were calculated at the mouse level so that each mouse served as the biological replicate. Mouse-level proportions were compared between groups using nonparametric tests with correction for multiple testing where appropriate.

### Exhaustion and naïve T cell scores

To quantify T-cell state at the single-cell level, we used UCell, a rank-based gene-signature scoring method, to calculate exhaustion- and naive-associated scores. The exhaustion-associated signature consisted of *Pdcd1, Lag3, Havcr2, Tigit, Ctla4, Nr4a1, Nr4a2, Nr4a3, Batf, Maf, Icos*, and *Eomes*, whereas the naive-associated signature consisted of *Lef1, Ccr7, Sell, Tcf7, Il7r, Klf2, S1pr1, Bach2, Foxo1*, and *Satb1*. In supplementary analyses, we additionally derived a net score by subtracting the naive score from the exhaustion score. For the main manuscript, exhaustion burden was summarized primarily using the exhaustion score across tissues, diet groups, and clonotype categories.

### TCR clonotype analysis

Matched TCR V(D)J annotations were integrated with single-cell transcriptomic metadata using the clonotype assignment associated with each T cell. For descriptive analyses, clone size was defined as the number of cells sharing the same clonotype and grouped into discrete categories. For the exhaustion-versus-clonotype analyses, clone size was recalculated locally within each mouse and organ to avoid inflating expansion through pooling across mice or tissues. Local clone categories were then summarized as single, small, and expanded, and exhaustion scores were compared across these categories. Correlations between clone size and exhaustion burden were assessed using Spearman correlation.

### Cell–cell communication analysis

Putative macrophage–T cell ligand–receptor communication was inferred using CellChat. Macrophage and T-cell populations from the relevant diet groups were analyzed after equalizing cell numbers across conditions. Communication probabilities were inferred using the mouse ligand–receptor reference database and were summarized at both the interaction and pathway levels. Comparative analyses focused on macrophage-derived signaling to T-cell populations, including exhausted T cells, across VD, HFD, and RD conditions. Because cells were pooled by condition rather than modeled as mouse-level replicates, these analyses were interpreted as predicted comparative communication networks rather than direct replicate-aware statistical evidence of signaling.

### Neutral-lipid staining

To assess tissue lipid accumulation, spleen, aorta, and lymph node samples were fixed, embedded, cryo-sectioned, and stained with BODIPY 493/503 and Nile Red, with DAPI used as a nuclear counterstain. Images were acquired under matched microscope settings across experimental groups. These studies were used to visualize diet-associated neutral-lipid accumulation in the tissue microenvironment.

### Flow-cytometric validation of splenic checkpoint expression

To validate exhaustion-associated programs at the protein level, splenic single-cell suspensions were prepared from mice at the late dietary time point and analyzed by multiparameter flow cytometry. After exclusion of debris, doublets, and dead cells, T cells were identified as CD45+CD3+ lymphocytes and then subdivided into CD4+ and CD8+ compartments. Surface expression of PD-1, TIGIT, LAG-3, TIM-3, and CTLA-4 was quantified using matched gating across groups. In addition to single-marker frequencies, co-expression of inhibitory receptors was measured to define multi-checkpoint-positive subsets, including PD-1+TIGIT+, PD-1+LAG-3+, and PD-1+TIM-3+ cells. Flow-cytometry results were summarized using mouse-level values as biological replicates.

### Statistical analysis

For mouse-level summaries, including composition analyses and flow-cytometric comparisons, the mouse was treated as the biological unit of replication. Two-group comparisons were generally performed using nonparametric tests, and correction for multiple testing was applied where appropriate. Correlations between clone size and exhaustion score were assessed using Spearman correlation. Data were visualized using standard plotting workflows with matched scaling and annotation across experimental groups.

## Results

### HFD reshapes T-cell composition across vascular and lymphoid tissues

To investigate how chronic dietary lipid exposure modulates adaptive immunity, we performed multi-tissue single-cell RNA sequencing (scRNA-seq) on CD45+ immune cells isolated from ApoE-/-mice at baseline and after 12 or 24 weeks on either a vegetarian diet (VD) or a high-fat diet (HFD), profiling the aorta, spleen, lymph nodes, bone marrow, and blood (**Fig. 1a**). Unsupervised clustering of the T-cell compartment identified seven distinct states, including naive/central memory, effector memory/effector, exhausted, regulatory, γδ, helper, and progenitor/proliferating T cells (**Fig. 1b-d**). Quantification of these populations revealed a clear diet-dependent redistribution of T-cell states, characterized by progressive contraction of the naive/central memory pool together with expansion of the exhausted T-cell population under HFD conditions (**Fig. 1c**). These compositional shifts were evident by 12 weeks and became more pronounced by 24 weeks, indicating that chronic lipid exposure drives sustained remodeling of T-cell differentiation.

**Figure 1.**
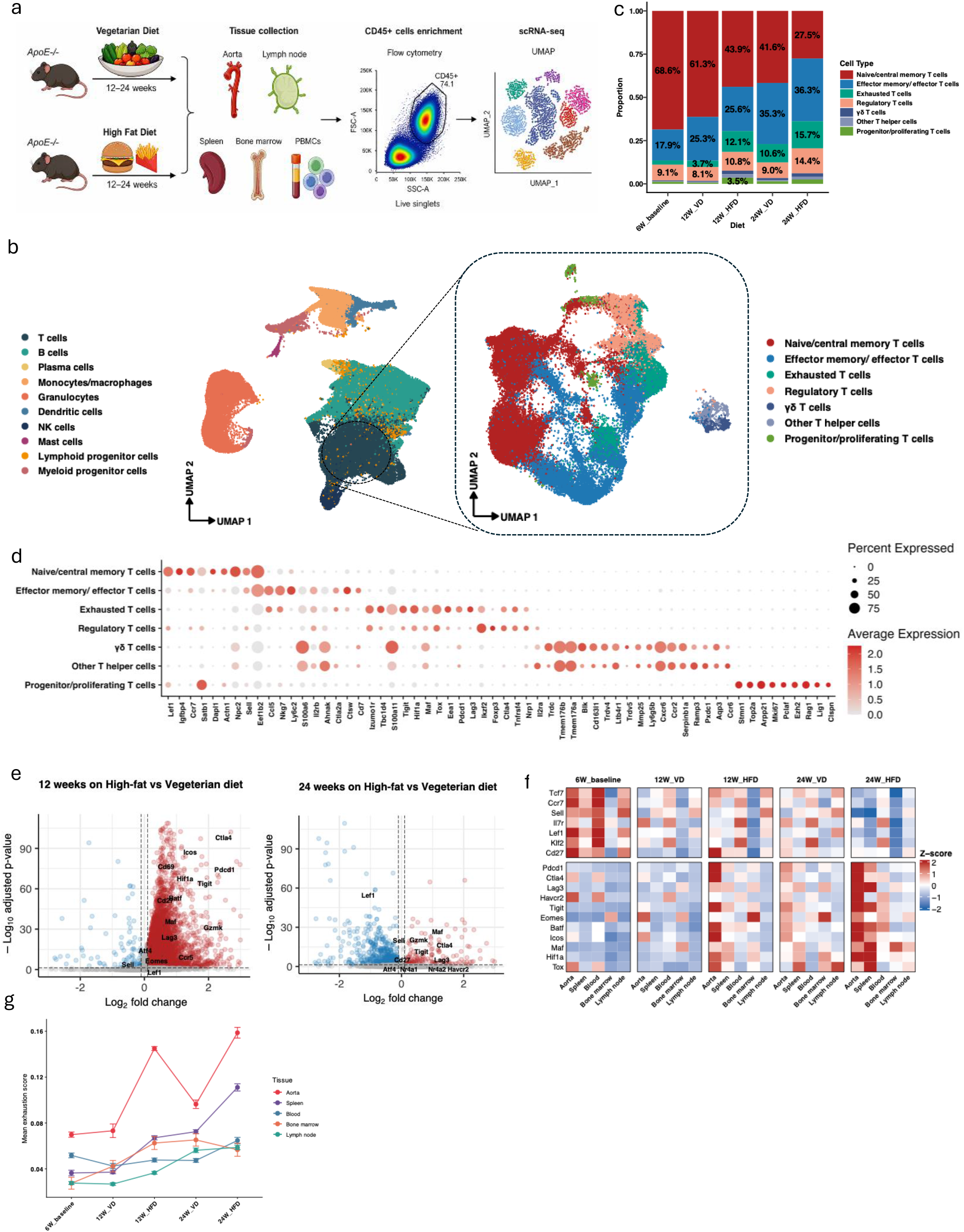
Chronic high-fat diet remodels T-cell composition and exhaustion-associated transcriptional programs across vascular and lymphoid tissues. a, Experimental workflow. ApoE□/□ mice were analyzed at baseline (6 weeks of age) or after 12 or 24 weeks of vegetarian control diet (VD) or high-fat diet (HFD). CD45□ immune cells isolated from the aorta, spleen, lymph nodes, bone marrow, and peripheral blood were profiled by single-cell RNA sequencing (scRNA-seq). b, Uniform manifold approximation and projection (UMAP) of all profiled immune cells, annotated by major cell population (left), and reclustered T cells, annotated by seven broad T-cell states (right). c, Relative proportions of annotated T-cell states among pooled T cells in each experimental group. Percentages indicate the contribution of selected populations; bars are descriptive pooled-cell summaries rather than mouse-level estimates. d, Expression of representative genes used to annotate T-cell states. Dot size indicates the percentage of cells expressing each gene, and color indicates average expression. e, Differential gene expression in T cells from HFD-versus VD-fed mice after 12 weeks (left) and 24 weeks (right). Positive log□ fold changes indicate higher expression under HFD. Selected genes are labeled; dashed lines indicate [specify fold-change and adjusted P-value thresholds]. f, Heatmaps of naive/memory-associated and activation/exhaustion-associated gene expression across tissues and experimental groups. Colors indicate Z-scored expression [specify aggregation and scaling procedure]. g, Mean exhaustion-associated UCell scores across tissues and experimental groups.

Differential expression analysis further supported this phenotypic shift. T cells from HFD-fed mice upregulated multiple genes associated with chronic activation and exhaustion-associated remodeling, including *Pdcd1* (PD-1), *Ctla4, Tigit, Lag3*, and *Havcr2* (TIM-3), whereas genes linked to less differentiated or memory-like states, including *Sell* and *Lef1*, were reduced (**Fig. 1e**). This transcriptional pattern was reproducible in an independent scRNA-seq dataset from a PCSK9-overexpressing hyperlipidemic mouse model (GSE239591), which showed increased expression of exhaustion-associated markers under HFD (**Supp. Fig. 1a, b**), supporting the generalizability of lipid-driven T-cell remodeling across distinct hyperlipidemic settings. Neutral-lipid staining confirmed lipid accumulation in vascular and lymphoid tissue sections, with more prominent BODIPY-positive deposits in HFD samples (**Supp Fig. 1c)**. To confirm these findings at the protein level, we performed flow cytometry on CD4+ and CD8+ T cells after 24 weeks of HFD and VD. Consistent with the scRNA-seq data, HFD-fed mice exhibited higher frequencies of inhibitory receptor-positive cells, including PD-1+, TIGIT+, TIM-3+, and LAG-3+ populations, as well as multi-inhibitory receptor-positive subsets, relative to VD controls (**Supp. Fig. 1d–g**).

We next examined whether this exhaustion-associated program was uniformly distributed across tissues. Although exhaustion-associated burden increased across all surveyed compartments under HFD, the magnitude of this increase was most pronounced in the aorta (**Fig. 1f,g; Supp. Fig. 1b**). By 24 weeks, aortic T cells exhibited the highest exhaustion scores and the lowest naive scores relative to lymphoid organs, identifying the aorta as the dominant site of diet-associated T-cell dysfunction. Collectively, these data show that chronic dietary lipid exposure broadly reshapes T-cell state across vascular and lymphoid tissues while driving its most pronounced exhaustion-associated remodeling within the aorta.

### Dietary reversal partially attenuates, but does not fully reset, exhaustion-associated T-cell remodeling

To determine whether HFD-driven T-cell remodeling is reversible, we analyzed a dietary reversal (RD) cohort in which ApoE-/-mice were maintained on HFD for 12 weeks before being switched to a vegetarian diet for an additional 12 weeks (**Fig. 2a**). At the global level, dietary reversal led to a partial restoration of the T-cell landscape, characterized by a reduction in the proportion of exhausted T cells and a relative recovery of the naive/central memory pool compared with mice maintained on sustained HFD (**Fig. 2b**). However, these shifts were incomplete, as the cell-state distribution in RD mice did not fully return to that observed in age-matched VD controls.

**Figure 2.**
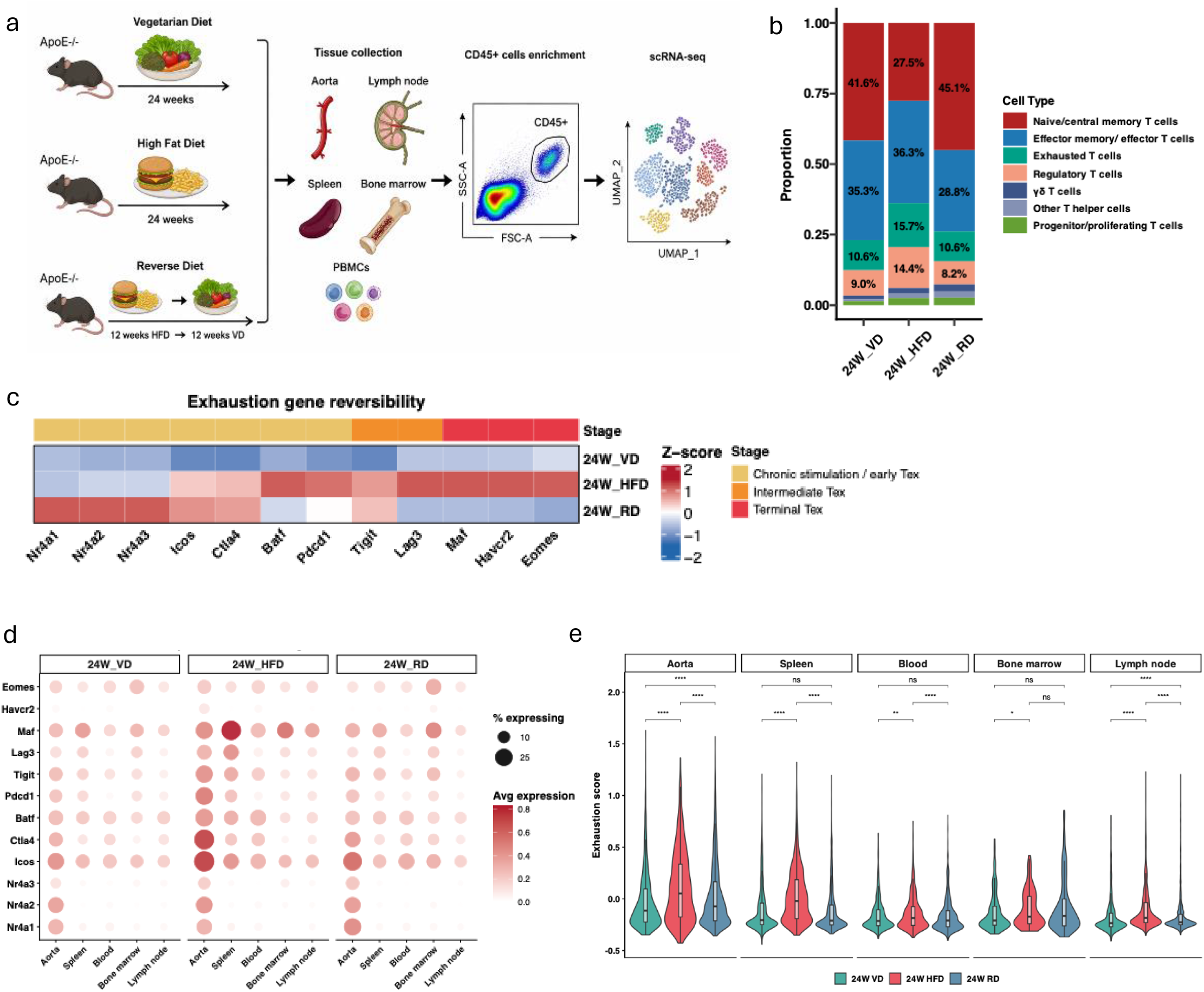
Dietary reversal alters T-cell composition and attenuates exhaustion-associated transcriptional programs in a tissue-dependent manner. a, Experimental design. ApoE□/□ mice received VD or HFD for 24 weeks, or HFD for 12 weeks followed by VD for 12 weeks (reversed diet, RD). CD45□ immune cells from five tissues were subjected to scRNA-seq. b, Relative proportions of annotated T-cell states among pooled cells in the 24-week VD, HFD, and RD groups. Percentages denote selected populations. c, Heatmap of selected activation-and exhaustion-associated genes across dietary conditions. Colors indicate Z-scored expression. The upper annotation groups genes into the indicated chronic-stimulation/early, intermediate, and terminal exhaustion-associated categories. d, Tissue-resolved expression of the genes shown in c. Dot size indicates the percentage of expressing cells, and color indicates average expression. e, Distributions of T-cell exhaustion-associated scores in the aorta, spleen, blood, bone marrow, and lymph nodes under VD, HFD, and RD conditions. Violins show score distributions; inset box plots show ns, not significant; *P < 0.05, **P < 0.01, ***P < 0.001, ****P < 0.0001 [P values are adjusted].

Gene-level analysis indicated that this partial recovery was accompanied by selective attenuation of exhaustion-associated transcriptional programs. Following dietary reversal, T cells showed reduced expression of genes associated with more advanced dysfunction and metabolic stress, including *Eomes, Pdcd1* (PD-1), *Havcr2* (TIM-3), *Lag3*, and *Hif1a* (**Fig. 2c**). In contrast, genes linked to early activation and chronic stimulation, most notably the Nr4a family (*Nr4a1, Nr4a2*, and *Nr4a3*), as well as *Icos* and *Tigit*, remained elevated in RD mice relative to VD controls (**Fig. 2c,d**). These findings suggest that removal of the dietary stimulus attenuates later exhaustion-associated programs but does not fully erase the earlier molecular imprint of chronic stimulation.

This pattern of incomplete recovery was consistent across tissues. Exhaustion scores were reduced by dietary reversal in all surveyed compartments but failed to fully normalize to VD levels in most tissues (**Fig. 2e**). Notably, the aorta, which exhibited the highest exhaustion burden under chronic HFD, remained the site with the most prominent residual exhaustion signal after reversal. Together, these findings indicate that lowering dietary lipid exposure partially restores the T-cell landscape and restrains progression of the exhaustion-associated program. However, because dietary reversal was initiated after 12 weeks of HFD exposure, these data are more consistent with attenuation of ongoing progression toward terminal dysfunction than with definitive reversal of a fixed, terminally exhausted state.

### Chronic HFD remodels the aortic macrophage–T cell niche toward T-cell activation, clonal expansion, and exhaustion

Because the aorta exhibited the strongest exhaustion phenotype under HFD, we next focused on the local aortic immune microenvironment to investigate programs and interactions associated with T-cell dysfunction. Differential expression analysis of aortic macrophages from HFD-fed mice revealed upregulation of inflammatory and antigen-presentation-associated genes, including MHC-related transcripts such as *H2-Ab1* and *H2-Eb1, Cd74*, interferon-responsive genes including *Stat1, Gbp2*, and *Batf2*, and chemokine mediators such as *Cxcl9* and *Cxcl10* (**Fig. 3a**). Pathway analysis further supported these findings, demonstrating enrichment of cytokine signaling, interleukin signaling, and antigen processing and presentation pathways in HFD-associated macrophages (**Fig. 3b**). Together, these data indicate that chronic lipid exposure shifts the aortic macrophage compartment toward an inflammatory and antigen-presenting state capable of supporting persistent T-cell stimulation.

**Figure 3.**
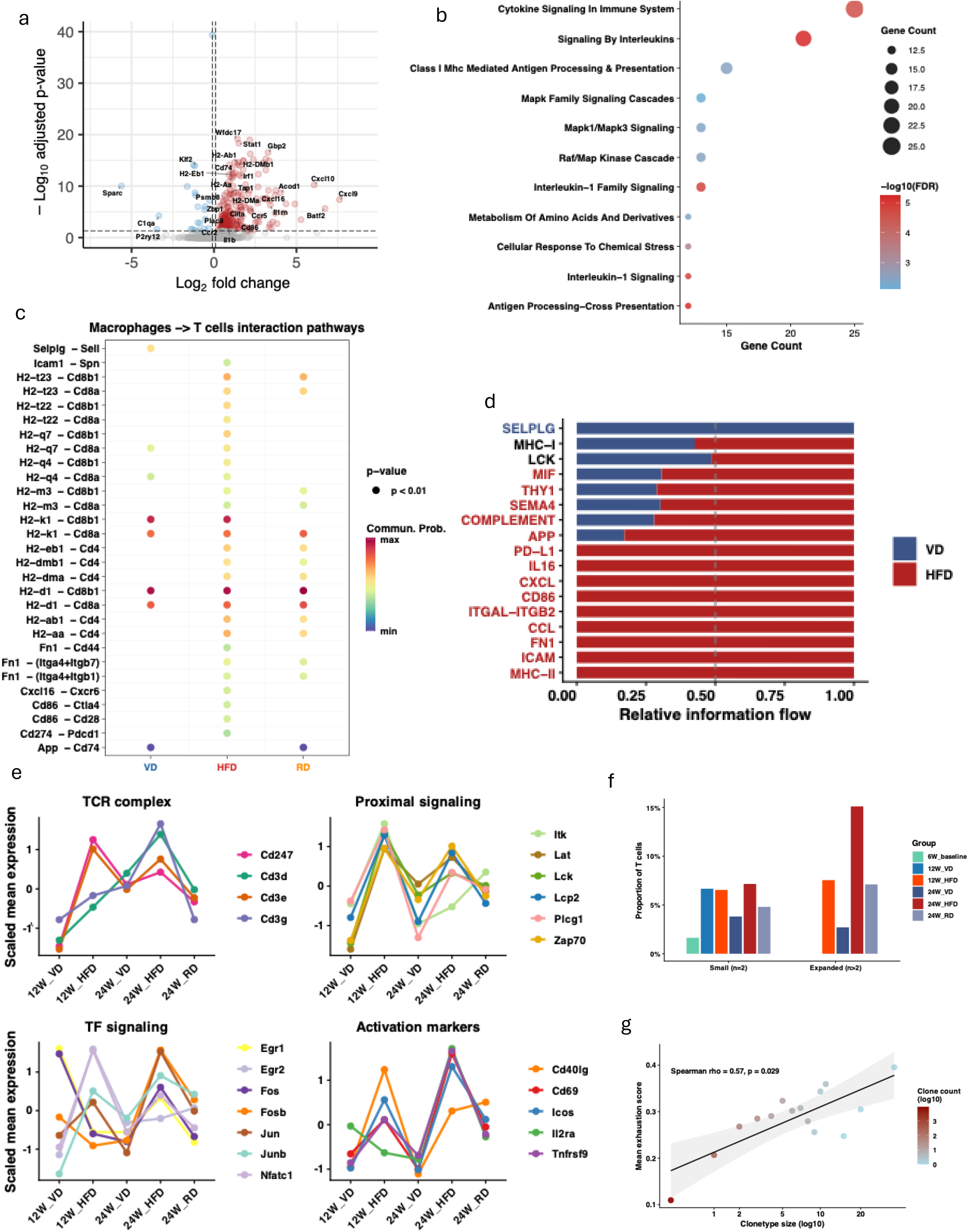
Aortic macrophage antigen-presentation programs and predicted macrophage– T-cell communication accompany T-cell activation and clonal expansion under HFD. a, Differential gene expression in aortic macrophages from HFD-versus VD-fed mice. Positive log□ fold changes indicate higher expression under HFD. Selected inflammatory, chemokine, and antigen-presentation-associated genes are labeled. b, Pathway enrichment analysis from the macrophage comparison in a. Dot size indicates the number of associated genes, and color indicates −log□□ false discovery rate (FDR). c, CellChat-inferred macrophage-to-T-cell ligand– receptor interactions across VD, HFD, and RD conditions. Dot color indicates inferred communication probability; displayed interactions meet the indicated CellChat P < 0.01 criterion. d, Relative information flow for the indicated signaling pathways in VD and HFD networks. Blue and red segments indicate the relative contributions of VD and HFD, respectively. e, Scaled mean expression of genes associated with the T-cell receptor (TCR) complex, proximal TCR signaling, transcription-factor responses, and activation in aortic T cells across diet groups and time points. f, Proportions of aortic T cells assigned to small (two cells) or expanded (more than two cells) clonotypes across experimental groups. g, Association between clonotype size and mean exhaustion-associated score. Clone sizes were calculated within each mouse and organ. Spearman ρ = 0.57; P = 0.029. For c, d, cell numbers were equalized across conditions before CellChat analysis. Cells were pooled by condition; inferred networks and interaction P values do not constitute mouse-level statistical evidence of altered signaling.

To determine how these macrophage changes might relate to the aortic T-cell phenotype, we performed CellChat analysis to infer macrophage-to-T-cell communication across VD, HFD, and RD conditions. HFD macrophages exhibited increased predicted outgoing communication through multiple pathways relevant to T-cell activation and dysfunction, including MHC-I and MHC-II, chemokine axes including CCL and CXCL, and costimulatory or checkpoint-associated signals such as CD86 and PD-L1 (**Fig. 3c,d**). Several of these pathways remained elevated in RD mice relative to VD controls, consistent with the incomplete recovery observed in the T-cell compartment following dietary reversal. These findings support a model in which chronic HFD amplifies macrophage-derived cues that sustain T-cell activation and promote acquisition of exhaustion-associated programs within the vascular wall.

Consistent with this heightened stimulatory environment, aortic T cells from HFD-fed mice showed coordinated upregulation of transcriptional programs linked to the TCR complex, including *Cd247, Cd3d, Cd3e*, and *Cd3g*; proximal signaling machinery, including *Lck, Zap70, Lat, Lcp2, Plcg1*, and *Itk*; immediate-early response factors, including *Egr1, Egr2, Fos, Fosb, Jun, Junb*, and *Nfatc1*; and activation markers such as *Cd69, Icos, Il2ra, Tnfrsf9*, and *Cd40lg* (Fig. 3e). These changes were accompanied by remodeling of clonal architecture, with chronic HFD increasing the proportion of expanded T-cell clonotypes, particularly at the 24-week time point (**Fig. 3f**). Moreover, expanded clonotypes exhibited higher exhaustion scores than singleton or small clones, and mean exhaustion score correlated positively with clonotype size (Spearman ρ = 0.57, p = 0.029; **Fig. 3g**). Collectively, these data indicate that chronic HFD is associated with enhanced aortic macrophage–T cell crosstalk, sustained T-cell activation, and clonal expansion, together with progressive acquisition of an exhaustion-associated dysfunctional phenotype.

## Discussion

Chronic dietary lipid exposure was associated with coordinated remodeling of T-cell states across vascular and lymphoid compartments, with the strongest exhaustion-associated transcriptional signal in the aorta. Dietary normalization reduced several HFD-associated changes but left selected activation-associated programs elevated, particularly within the vascular compartment. These observations were accompanied by inflammatory and antigen-presentation programs in aortic macrophages, increased predicted macrophage-to-T-cell communication, and an association between T-cell clonal expansion and exhaustion scores.

The multi-tissue design places vascular T-cell remodeling within a broader systemic response. Human plaque profiling has previously identified activated, differentiated, and exhaustion-associated T-cell populations that differ from circulating T cells.^8,12^ The present study extends this framework by comparing vascular, lymphoid, and circulating compartments at defined stages of dietary exposure. Although the magnitude and temporal pattern of the response differed among tissues, the aorta consistently showed the greatest exhaustion-associated burden. This distribution suggests that systemic dietary exposure alone does not determine T-cell state; the local vascular environment may amplify or sustain responses initiated elsewhere. Whole-aorta profiling, however, cannot assign these changes exclusively to intimal plaques or distinguish them from responses in the adventitia and other vascular niches.

The coexistence of enhanced activation and exhaustion-associated programs is consistent with the context-dependent effects of cholesterol on T cells. Hypercholesterolemia can enhance TCR signaling and responsiveness to stimulation, whereas cholesterol accumulation in tumor-infiltrating CD8+ T cells can induce endoplasmic reticulum stress and XBP1-dependent inhibitory receptor expression^5,6^The present findings suggest that heightened responsiveness and subsequent dysfunction need not represent contradictory outcomes. Instead, persistent stimulation in a lipid-rich inflammatory environment may favor both activation and acquisition of regulatory programs that constrain sustained responses. Nevertheless, the mouse experiments do not isolate cholesterol from other dietary differences, and neither T-cell cholesterol loading nor the ER stress–XBP1 pathway was directly established as the mechanism underlying the observed phenotype.

Dietary reversal revealed an important distinction between recovery of population composition and normalization of transcriptional state. The pooled proportion of cells annotated as exhausted returned to the VD value, while selected chronic-stimulation-associated transcripts remained elevated and residual differences were tissue dependent. Thus, a restored population frequency does not necessarily indicate complete molecular recovery. Conversely, persistent expression of NR4A-family genes or inhibitory receptors does not establish irreversible exhaustion. Such persistence could reflect ongoing antigen exposure, residual tissue inflammation, altered trafficking, or replacement of one T-cell population by another. Because dietary reversal began after 12 weeks of HFD and individual cells were not tracked, the data cannot distinguish reprogramming of previously affected cells from prevention of further remodeling or changes in population turnover. Durable epigenetic changes are recognized in established exhaustion, but an analogous epigenetic imprint remains to be demonstrated in this model^13^.

The macrophage analyses identify a plausible cellular link between chronic dietary exposure and vascular T-cell stimulation. HFD-associated macrophages expressed inflammatory, interferon-responsive, and antigen-presentation programs, while communication inference suggested increased capacity for antigen recognition, co-stimulation, chemotaxis, and checkpoint engagement. These findings are compatible with an environment that supports repeated T-cell encounters rather than uniformly suppressing or activating immunity. Related mechanisms have been demonstrated in cancer, where macrophage antigen presentation and persistent macrophage–T-cell interactions can promote exhaustion^14,15^. This comparison provides a mechanistic precedent, not evidence that the same pathway operates identically in atherosclerosis. CellChat predictions remain hypothesis generating, particularly because the networks were inferred from pooled conditions rather than mouse-level replicates; they do not establish physical contact, ligand activity, or macrophage necessity in vivo.

The association between clonal expansion and exhaustion scores provides another link between repeated stimulation and T-cell remodeling. Larger locally defined clonotypes exhibited greater exhaustion-associated burden, consistent with selective expansion or persistence of antigen-experienced populations. APOB-specific T-cell responses have been demonstrated in humans, and their magnitude and phenotype can vary with hyperlipidemia and coronary disease .^12^.Nevertheless, clonotype expansion in the present dataset does not identify antigen specificity, establish where expansion occurred, or demonstrate that expansion preceded exhaustion-associated remodeling. Direct antigen mapping would be needed to distinguish responses to atherosclerosis-associated self-antigens from unrelated memory or bystander populations.

An important implication is that exhaustion-associated features should not automatically be interpreted as pathogenic or therapeutically undesirable. In atherosclerosis, inhibitory receptors can restrain damaging immune responses, while some checkpoint-positive T cells retain substantial inflammatory activity. Mulholland et al. identified progenitor-exhausted PD-1-positive aortic T cells capable of producing IFN-γ and showed that PD-1 blockade increased vascular T-cell activation and accumulation.^11^ The present broad annotations do not resolve equivalent progenitor, intermediate, and terminal states. Consequently, a higher exhaustion score cannot be equated with either greater plaque injury or effective immune protection. The findings support investigating how dietary and antigen-presenting-cell signals alter the balance between persistent inflammation and immune restraint, rather than assuming that generalized reversal of exhaustion would improve vascular outcomes.

Several limitations constrain the conclusions. The predominantly female ApoE-deficient mouse model requires validation across sexes and additional disease settings. The diets differed in energy density and composition, preventing attribution of all effects specifically to cholesterol. Tissue collection at successive endpoints provides a time-course comparison, not longitudinal tracking of the same cells or animals. Relative cell proportions are sensitive to changes in other populations and do not establish absolute expansion or depletion. In addition, the exhaustion signature includes genes shared with activation and other T-cell states, and aggregate scores may reflect both compositional shifts and changes within subsets. Establishing stable exhaustion will require subset-resolved functional testing, recovery after stimulus withdrawal, and epigenetic characterization, rather than inhibitory receptor expression alone.^13^

Overall, these findings connect chronic HFD exposure with systemic T-cell remodeling and a particularly prominent vascular exhaustion-associated phenotype. Dietary normalization modifies this response without uniformly restoring all transcriptional features, while macrophage conditioning and repeated-stimulation experiments support a potential antigen-presentation-dependent contribution to exhaustion-like differentiation. The resulting framework links dietary exposure, local macrophage state, and persistent T-cell stimulation while distinguishing those associations from proven in vivo causality. Determining how these processes influence plaque burden and stability will be essential before their modulation can be considered a strategy for vascular protection

## Conclusion

Chronic high-fat diet promotes progressive T cell state remodeling across vascular and lymphoid tissues during atherogenesis, characterized by lipid accumulation, reduced naive/memory-associated features, increased exhaustion-like and inhibitory receptor-associated programs. These findings suggest that dietary lipid exposure may leave a durable adaptive immune imprint across organs during atherosclerosis.

## Figure Legends

**Supplementary Figure 1.**
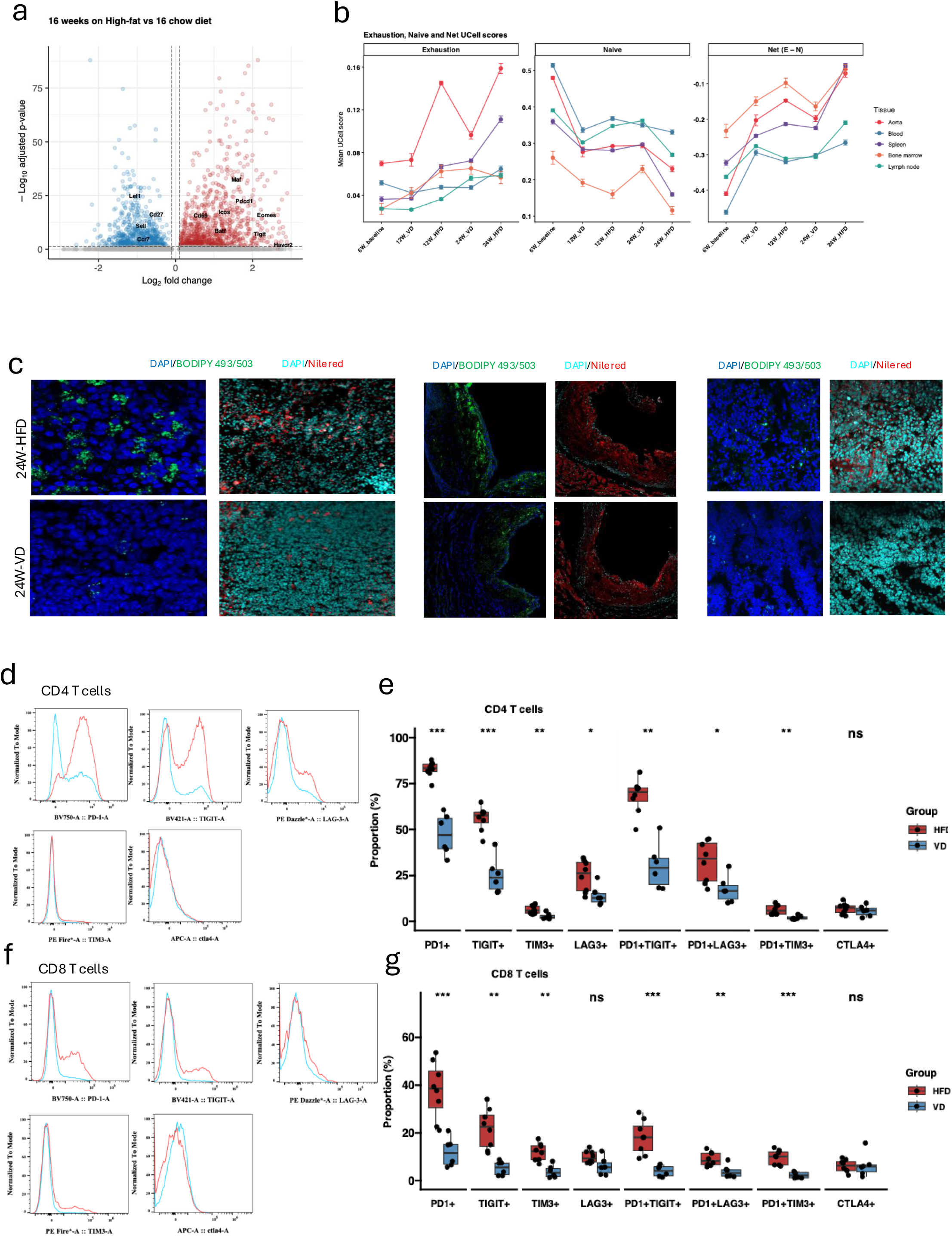
Independent transcriptomic analysis, tissue lipid staining, and splenic inhibitory-receptor phenotyping complement the multi-tissue T-cell analysis. a, Differential gene expression in T cells from an independent PCSK9-overexpression hyperlipidemia dataset (GSE239591), comparing 16 weeks of HFD with chow diet. Positive log□ fold changes indicate higher expression under HFD. Selected genes are labeled; dashed lines indicate [specify thresholds]. b, Mean exhaustion-associated, naive-associated, and net UCell scores across tissues, time points, and diet groups. The net score was calculated as the exhaustion-associated score minus the naive-associated score. c, Representative fluorescence images of spleen after 24 weeks of HFD or VD. BODIPY 493/503 and Nile Red were used to visualize lipid staining, with DAPI counterstaining of nuclei. Images were acquired under matched settings across diet groups. d-g. Representative flow-cytometry histograms of PD-1, TIGIT, LAG-3, TIM-3, and CTLA-4 expression in splenic CD4□ (d) and CD8□ (f) T cells after 24 weeks of HFD or VD. Mouse-level frequencies of inhibitory-receptor-positive and dual-positive subsets among splenic CD4□ (e) and CD8□ (g) T cells. Red denotes HFD and blue denotes VD. Each point represents one mouse; ns, not significant; *P < 0.05, **P < 0.01, ***P < 0.001 [P values are adjusted].

## References

1. Dutta P, Sager HB, Stengel KR, et al. Myocardial Infarction Activates CCR2(+) Hematopoietic Stem and Progenitor Cells. Cell Stem Cell. May 07 2015;16(5):477–87. doi:10.1016/j.stem.2015.04.008

2. Libby P, Buring JE, Badimon L, et al. Atherosclerosis. Nat Rev Dis Primers. Aug 16 2019;5(1):56. doi:10.1038/s41572-019-0106-z

3. Wolf D, Ley K. Immunity and Inflammation in Atherosclerosis. Circ Res. Jan 18 2019;124(2):315–327. doi:10.1161/CIRCRESAHA.118.313591

4. Libby P, Hansson GK. From Focal Lipid Storage to Systemic Inflammation: JACC Review Topic of the Week. J Am Coll Cardiol. Sep 24 2019;74(12):1594–1607. doi:10.1016/j.jacc.2019.07.061

5. Mailer RKW, Gisterå A, Polyzos KA, Ketelhuth DFJ, Hansson GK. Hypercholesterolemia Enhances T Cell Receptor Signaling and Increases the Regulatory T Cell Population. Sci Rep. Nov 15 2017;7(1):15655. doi:10.1038/s41598-017-15546-8

6. Ma X, Bi E, Lu Y, et al. Cholesterol Induces CD8. Cell Metab. Jul 02 2019;30(1):143–156.e5. doi:10.1016/j.cmet.2019.04.002

7. Saigusa R, Winkels H, Ley K. T Cell Subsets and Functions in Atherosclerosis. Nature Reviews Cardiology. 2020/07/01 2020;17(7):387–401. doi:10.1038/s41569-020-0352-5

8. Fernandez DM, Rahman AH, Fernandez NF, et al. Single-Cell Immune Landscape of Human Atherosclerotic Plaques. Nat Med. Oct 2019;25(10):1576–1588. doi:10.1038/s41591-019-0590-4

9. Depuydt MAC, Prange KHM, Slenders L, et al. Microanatomy of the Human Atherosclerotic Plaque by Single-Cell Transcriptomics. Circ Res. Nov 6 2020;127(11):1437–1455. doi:10.1161/circresaha.120.316770

10. Chowdhury RR, Valainis JR, Kask O, et al. The Role of Antigen Recognition in the γδ T Cell Response at the Controlled Stage of M. tuberculosis Infection. bioRxiv. 2021:2021.09. 14.460324.

11. Mulholland M, Chalou A, Andersson SHA, et al. Progenitor Exhausted PD-1. Nat Cardiovasc Res. Oct 2025;4(10):1311–1328. doi:10.1038/s44161-025-00713-2

12. Chowdhury RR, D’Addabbo J, Huang X, et al. Human Coronary Plaque T Cells Are Clonal and Cross-React to Virus and Self. Circ Res. May 13 2022;130(10):1510–1530. doi:10.1161/CIRCRESAHA.121.320090

13. Kang TG, Johnson JT, Zebley CC, Youngblood B. Epigenetic Regulation of T Cell Exhaustion in Cancer. Nat Rev Cancer. Jan 2026;26(1):46–61. doi:10.1038/s41568-025-00883-y

14. Kersten K, Hu KH, Combes AJ, et al. Spatiotemporal Co-Dependency between Macrophages and Exhausted CD8. Cancer Cell. Jun 13 2022;40(6):624–638.e9. doi:10.1016/j.ccell.2022.05.004

15. Nixon BG, Kuo F, Ji L, et al. Tumor-Associated Macrophages Expressing the Transcription Factor IRF8 Promote T cell Exhaustion in Cancer. Immunity. Nov 08 2022;55(11):2044–2058.e5. doi:10.1016/j.immuni.2022.10.002

